# Degeneracy measures in biologically plausible random Boolean networks

**DOI:** 10.1101/2021.04.29.441989

**Authors:** Basak Kocaoglu, William Alexander

**Affiliations:** Center for Complex Systems and Brain Sciences, Florida Atlantic University, Boca Raton, Florida, USA

## Abstract

Biological systems show diversity in terms of the underlying network structure and the governing rules of such networks. Yet, different types of biological networks may develop similar adaptation strategies in face of environmental changes. Degeneracy refers to the ability to compensate for compromised function without the need for a redundant component in the system. Highly degenerate systems show resilience to perturbations and damage because the system can compensate for compromised function due to reconfiguration of the underlying network dynamics.

Although formal definitions of degeneracy have been proposed, these definitions have only been tested in relatively simple networks involving weighted connections between network nodes. In this study, we test an information theoretic definition of degeneracy on random Boolean networks, frequently used to model gene regulatory networks. Random Boolean networks are discrete dynamical systems with binary connectivity and thus, these networks are well-suited for tracing information flow and the causal effects. By generating networks with random binary wiring diagrams, we test the effects of systematic lesioning of connections and perturbations of the network nodes on the degeneracy measure.

Our analysis shows that degeneracy, on average, is the highest in networks in which ~20% of the connections are lesioned while 50% of the nodes are perturbed. Moreover, our results for the networks with no lesions and the fully-lesioned networks are comparable to the degeneracy measures from weighted networks, thus we show that the degeneracy measure is applicable to different networks. Such a generalized applicability implies that degeneracy can be used to make predictions about the variety of systems’ ability to recover function.

**Author Summary:** Degeneracy – the ability of structurally different elements to perform similar functions – is a property of many biological systems. Systems exhibiting a high degree of degeneracy continue to exhibit the same macroscopic behavior following a lesion even though the underlying network dynamics are significantly different. Degeneracy thus suggests how biological systems can thrive despite changes to internal and external demands. Although degeneracy is a feature of network topologies and seems to be implicated in a wide variety of biological processes, research on degeneracy in biological networks is mostly limited to weighted networks (e.g., neural networks). To date, there has been no extensive investigation of information theoretic measures of degeneracy in other types of biological networks. In this paper, we apply existing approaches for quantifying degeneracy to random Boolean networks used for modeling biological gene regulatory networks. Using random Boolean networks with randomly generated rulesets to generate synthetic gene expression data sets, we systematically investigate the effect of network lesions on measures of degeneracy. Our results are comparable to measures of degeneracy using weighted networks, and this suggests that degeneracy measures may be a useful tool for investigating gene regulatory networks.

## Introduction

Biological systems can adjust their functioning dynamically in face of changing circumstances. However, such functional adjustments are constrained by the structural properties of the components that perform the functions [1–7] as well as the topology of the system. Biological systems as complex networks have evolved multiple strategies to achieve a ‘working’ reconfiguration of the components that ensures survival through shifts in environmental contingencies [8–15]. One strategy is redundancy which means that a system has multiple structurally identical components serving the same function [16–19]. Systems also can utilize multifunctionality (or alternatively, pluripotentiality [20,21]) that is the capacity for a single component to serve multiple functions [22–26]. Another strategy for biological systems to respond flexibly to perturbations is called degeneracy [9,10,27–29].

Degeneracy (or alternatively, ‘distributed redundancy’ [30], ‘distributed robustness’ [31], ‘functional redundancy’ [25], ‘extrinsic buffering’ [32]) describes the ability of components in a biological system that are structurally different to carry out the same or similar functions [20,21,21,28,33–40]. As difference in the structure implies different functions [41], under certain conditions degenerate components do not necessarily show functional variety [42,43] but instead, each degenerate component is responsible for a (set of) function(s) which is initially determined by their biochemical(/physical) structure [9,21,31]. Unlike multifunctionality and redundancy, degeneracy implies a change in the role assignments among the components such that the system can continue to function even when its normal processes have been compromised.

A biological network with high degeneracy means that the system can show the same macroscopic behavior following a lesion even though the underlying network dynamics are significantly different. In other words, if the system is highly degenerate, after a lesion, the function can be recovered by a structurally different (i.e., performing a different function under normal conditions) component taking over a new function. For example, in the brain many different neural clusters can affect the same motor outputs, and if some of the brain areas are damaged, an alternative (non-redundant) pathway can be recruited in order to generate functionally equivalent behaviors [44–48]. Degeneracy thus suggests how biological systems can thrive despite changes to internal and external demands.

It has been shown that degeneracy also plays a role in complexity and evolvability of the biological systems [8,10,27,28,32,36,49]. Higher levels of degeneracy correlate with an increase in the degree of both the functional integration and local segregation of a system, and therefore, higher degeneracy is accompanied by higher degree of complexity of the systems [33].While local segregation (namely, functional specialization) enables system to be flexible against environmental stress (due to diversity of functions), functional integration allows system to be robust [50–52]. If a component, or a group of components are compromised in a highly degenerate system, functions can be reassigned among distinct elements (that are locally segregated) while the macro-level behavior (which requires the system to be functionally integrated) is conserved. This adaptability brings an obvious advantage over the course of natural selection [44].

To measure degeneracy in systems, Tononi et al. [33] introduced a quantitative measure for neural networks (see also alternatives [53]) using an information theoretic approach. Information theory [54–56] provides a set of tools to describe how information is processed in systems. It allows us to measure the statistical (in)dependencies in terms of the information content of the components. Mutual information (MI), which is a measure provided by information theory, can also capture nonlinear dependencies(/relationships) that are not detectable by correlation analysis [54,57]. However, direction of the interaction between the components cannot be discerned from MI alone [33,57,58]. Incorporating MI with systematic perturbations (to determine directionality), degeneracy is formalized [33] in terms of the causal effects of the changes in the state of the subsets (components and/or subgroups of components) on the system’s output. If the output activity of the system is not affected by the change (e.g., perturbation) in a subset’s state, then the system is highly degenerate with regard to the function that is performed by that subset. This information theoretic measure of degeneracy is, first, applied to highly abstract networks in the work by Tononi et al. [33] which is followed by applications to the weighted networks with a high degree of biological fidelity (e.g., Hodgkin-Huxley type neural networks [40] and genetic networks with epistasis [59]).

Although it has been shown that degeneracy as a network property exists at different levels of biological organization (from molecules to behavior [30,36,44]), a quantitative analysis of degeneracy at such levels is sparse and methods are individualized to specific cases (see the different versions of degeneracy measurements in other works [10,53,60]). In systems biology, information theoretic measures are widely applied to many problems [61], yet, to date, there is not a comprehensive study applying this measure for biologically realistic networks other than networks with weighted connections.

Neural networks offer one example of how biological systems can incorporate degeneracy to ensure survival after being damaged. However, other biological networks are likewise capable of recovering partial or full function following damage. For example, in between-species interaction networks a species loss can be compensated by other species contributing to ecosystem functioning [62]. Likewise, on a smaller scale, it has been shown that loss of functioning in some (non-redundant) genes has a weak or no effect on the fitness of the gene networks [8]. Although degeneracy might not be detectable under normal conditions, after perturbating or lesioning the biological networks, changes in the environment may also evoke degenerate responses (‘degeneracy lifting’ effect [9]). Environmental (evolutionary) pressure in receptor-signal transduction networks [53] can push the signaling pathways to reconfigure into a degenerate form.

Although degeneracy is a feature of biological gene networks, it is unclear whether models of gene networks can be analyzed using the same information theoretic approach as used for neural network models. A GRN is a network of gene-gene interactions through their regulators that control the gene expression levels of the products (mRNAs and proteins) which, ultimately, determine the cell fate (final cell type, i.e., function of the cell) [63]. Measurements of degeneracy at the level of gene transcription control may provide insights on how functions of genes that determine the cell function, can be recovered as a consequence of the network properties (GRN topology).

Random Boolean network (RBN) models, as discrete models of GRNs, are well-suited to study degeneracy since with RBNs we can induce and trace the effects of targeted lesions while environmental/external pressure is a parameter that can be controlled over in silico experiments. Unlike neuronal networks where edges are (synaptic) weight vectors, RBNs have a static wiring diagrams [64] governed by logic equations that represent the functions of gene regulatory factors (e.g., transcription factors). Logic equations describe the underlying network architecture. For example, for a simple network of 3 genes, if gene G1 is regulated both directly by G3 and through an indirect link from G2, this architecture is represented by the logic function of “G1 AND (G2 OR G3)”. Likewise, if there is an inhibitory regulation of G1 through G2 while the same architecture is preserved from the network described above, this structure can be represented by the logic equation of “G1 AND ((NOT G2) OR G3)”. Since each state of gene expression is the direct outcome of the activity of (regulatory interactions in) the previous state, one can assess the effects of circuit architecture on gene expression levels [63]. Hence, in RBNs, it is feasible to trace the information flow at each (discrete) time step and so, causal influences.

In this study we test to what extent the information theoretic measure of degeneracy applies to RBNs. Furthermore, we test systematic lesions in randomly generated Boolean networks while varying the number of perturbed nodes. This enables us to explore how degeneracy quantitively changes as a function of interventions to the nodes and induced topological alterations in the networks. Results show that degeneracy measures are comparable to different networks – not only to weighted networks.

## Results

### Lesioning

We first investigate the effects of systematic lesioning on measures of degeneracy in randomly wired RBNs. Edges between network nodes were lesioned in two ways. In type-1 lesioning, only outgoing edges were lesioned incrementally while it is possible (due to the pseudo-random algorithm to generate the logic functions) that incoming edges stayed intact and for all the nodes, self-connections were preserved in the network (for details see Methods). The networks with type-1 lesions decrease in average degeneracy values as the cut percentage increases (Fig1). This validates that degeneracy emerges as a network property.

**Fig1.**
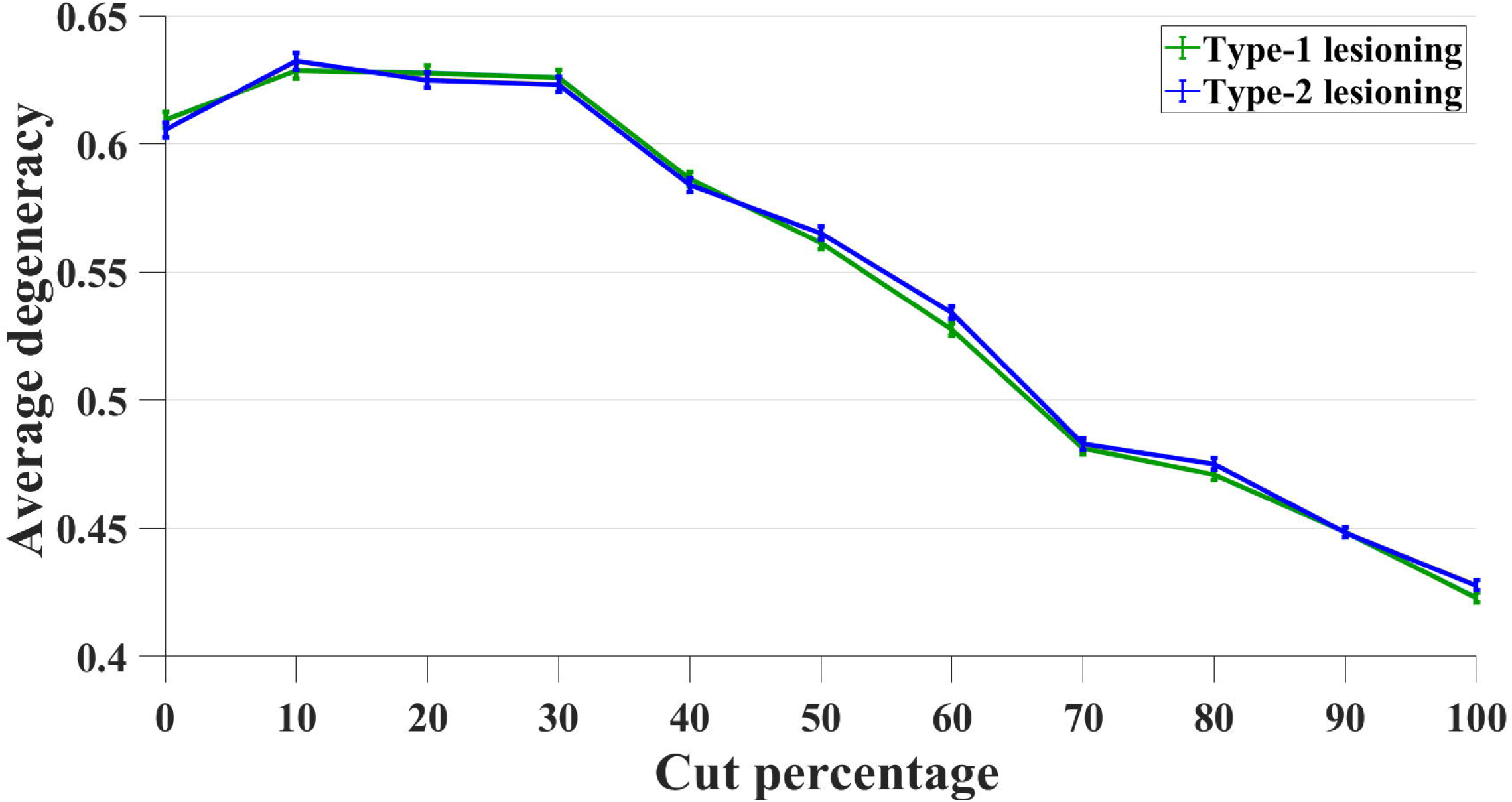
Average degeneracy values compared between type-1 (green line) and type-2 (blue line) lesioning. On the x axis, the cut percentage represents the affected number of nodes (in total of 10 nodes) whose edges are lesioned given the networks. For both lesioning types, degeneracy was lowest in the 100% cut condition where edges of all nodes (10) were cut.

In the second type of lesioning (hence the name type-2 lesion), we lesioned all incoming and outgoing edges (but not the self-connections) for randomly chosen genes incrementally. Similar to the type-1 lesioning, average degeneracy decreases as a function of lesioning. We have anticipated that there could be a difference between the effects of lesioning types as a direct consequence of the partial lesioning (the incoming edges are preserved) in type-1 condition which can lead some nodes to become dead ends since the activity ends in those nodes. Active nodes without outgoing edges means that such nodes do not serve a function, and this eventually would result in lower degeneracy values. The comparison of two lesioning types, in Fig 1, demonstrates that both conditions have similar effect on the average degeneracy, where there is a no significant difference (two-way analysis of variance, ANOVA) found between both types of lesioning. However, average degeneracy varies with cut conditions (p = 0.004).

### Perturbation

Degeneracy is calculated as the area between the average *MI^P^(X^k^;O)* (mutual information, MI, between the portion of entropy shared by the system for *each* perturbed subset k and the output *O*) and overall-MI (mutual information between the system and its output) for different perturbed subset sizes, k (see Equation 3 in Methods). This area shows a characteristic shape of the degeneracy function: a non-zero value that declines to zero as perturbed subset size k approaches *k = O*, following an increase that is “higher than would be expected from a linear increase” [33]. This characteristic shape has furthermore been replicated in networks composed of in Hodgkin-Huxley neurons [40]. In our study, the analogous condition for such illustration of degeneracy is where no edges are lesioned in the networks (Fig2a).

**Fig 2.**
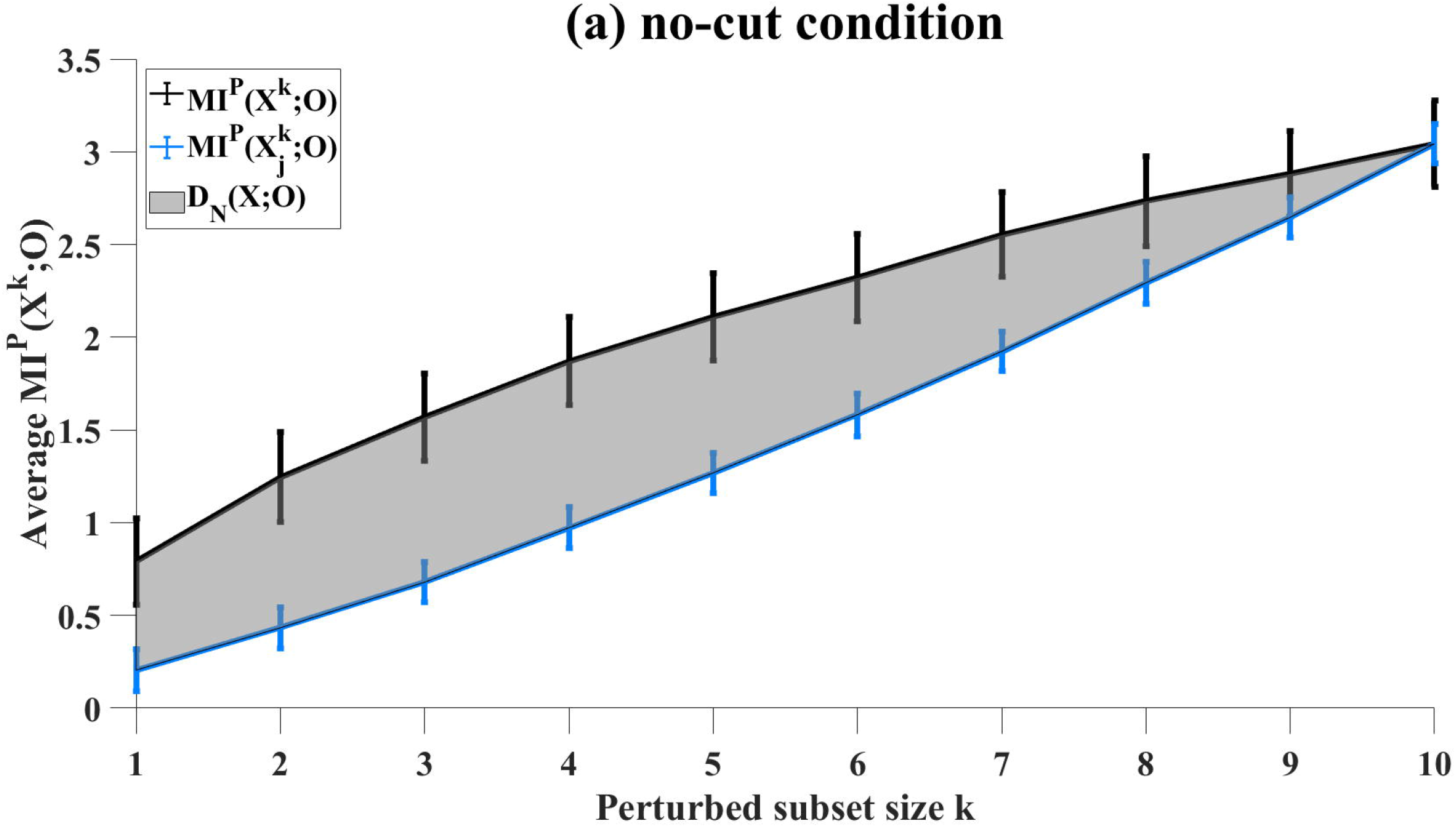
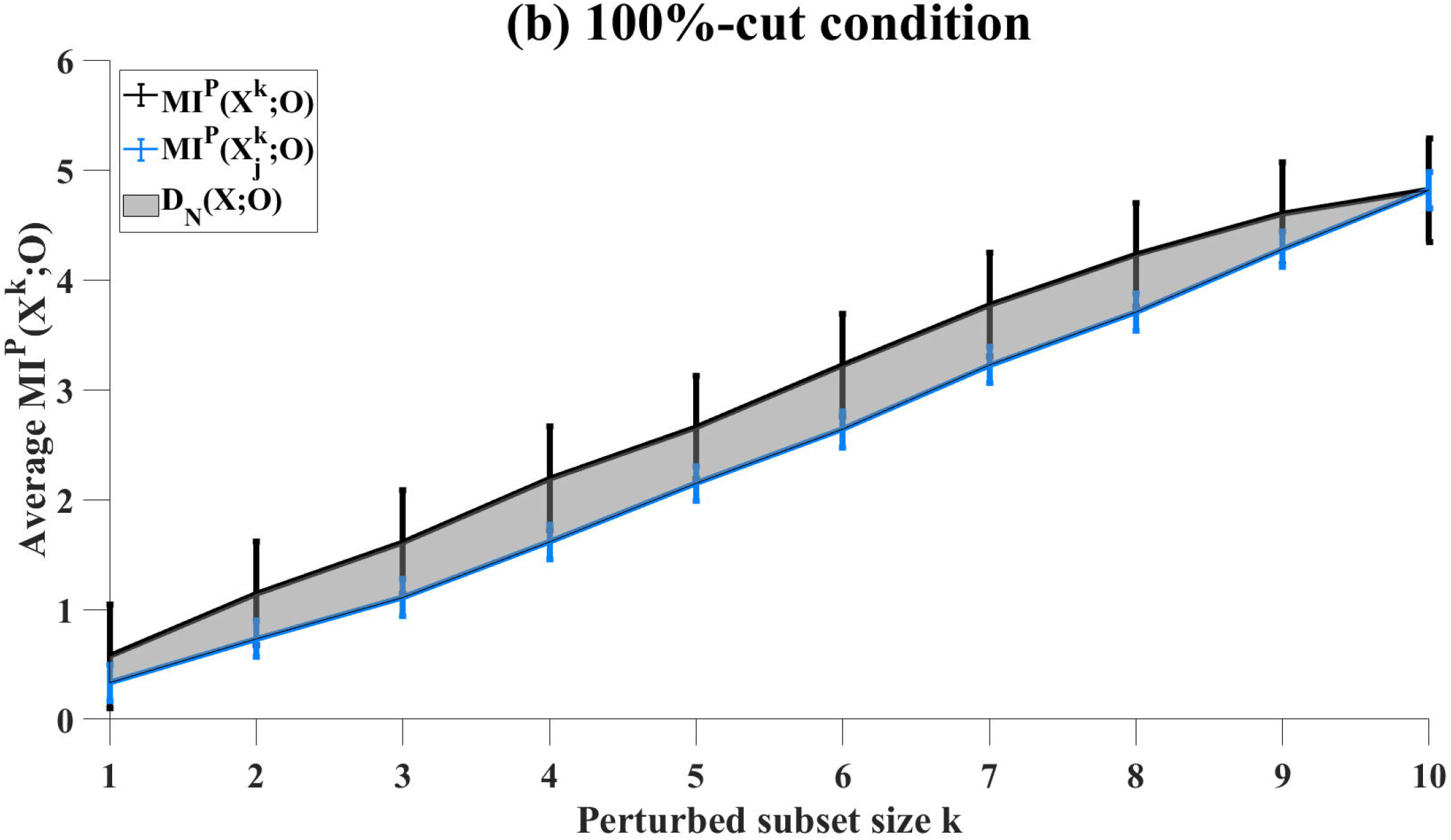
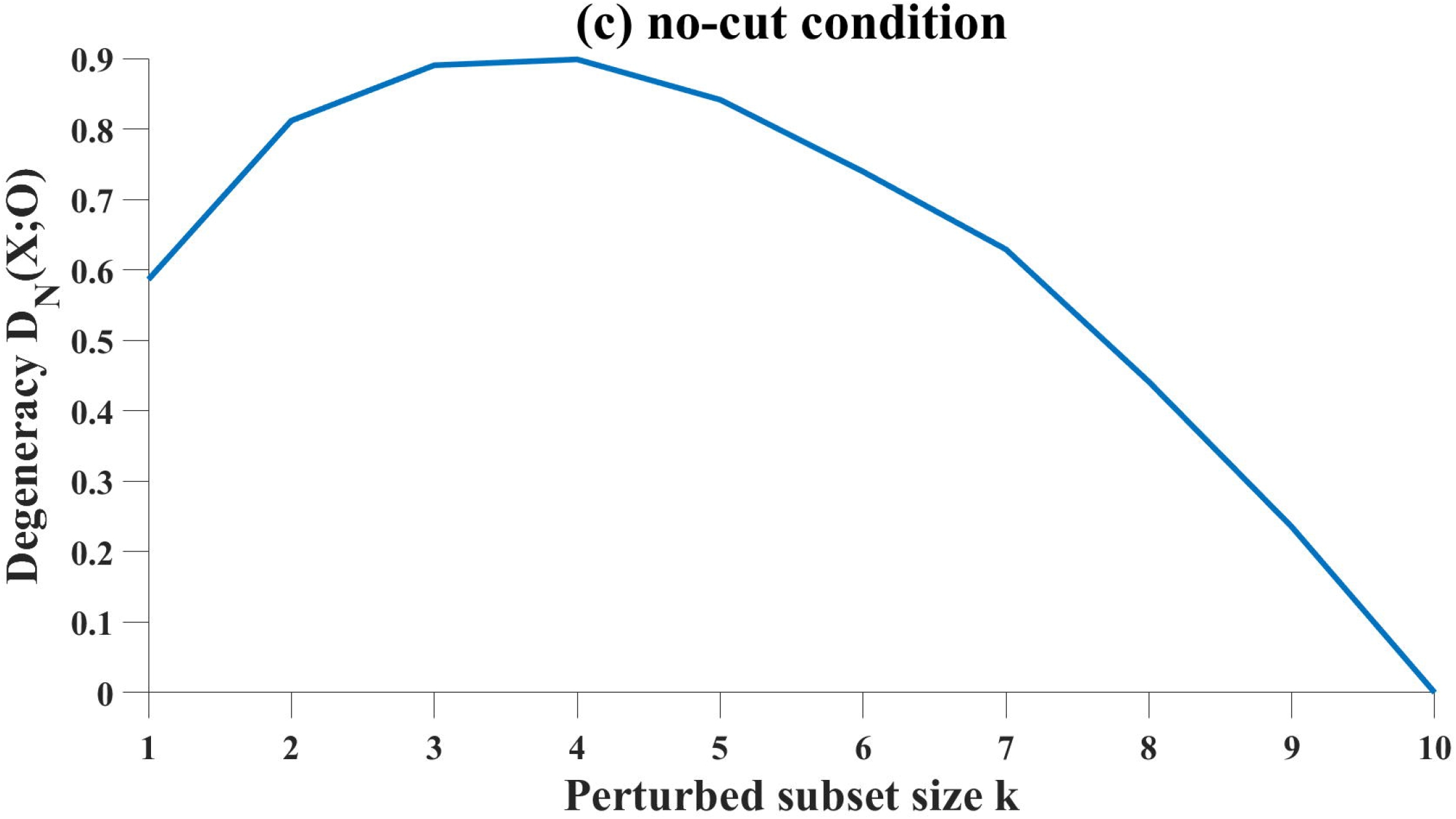
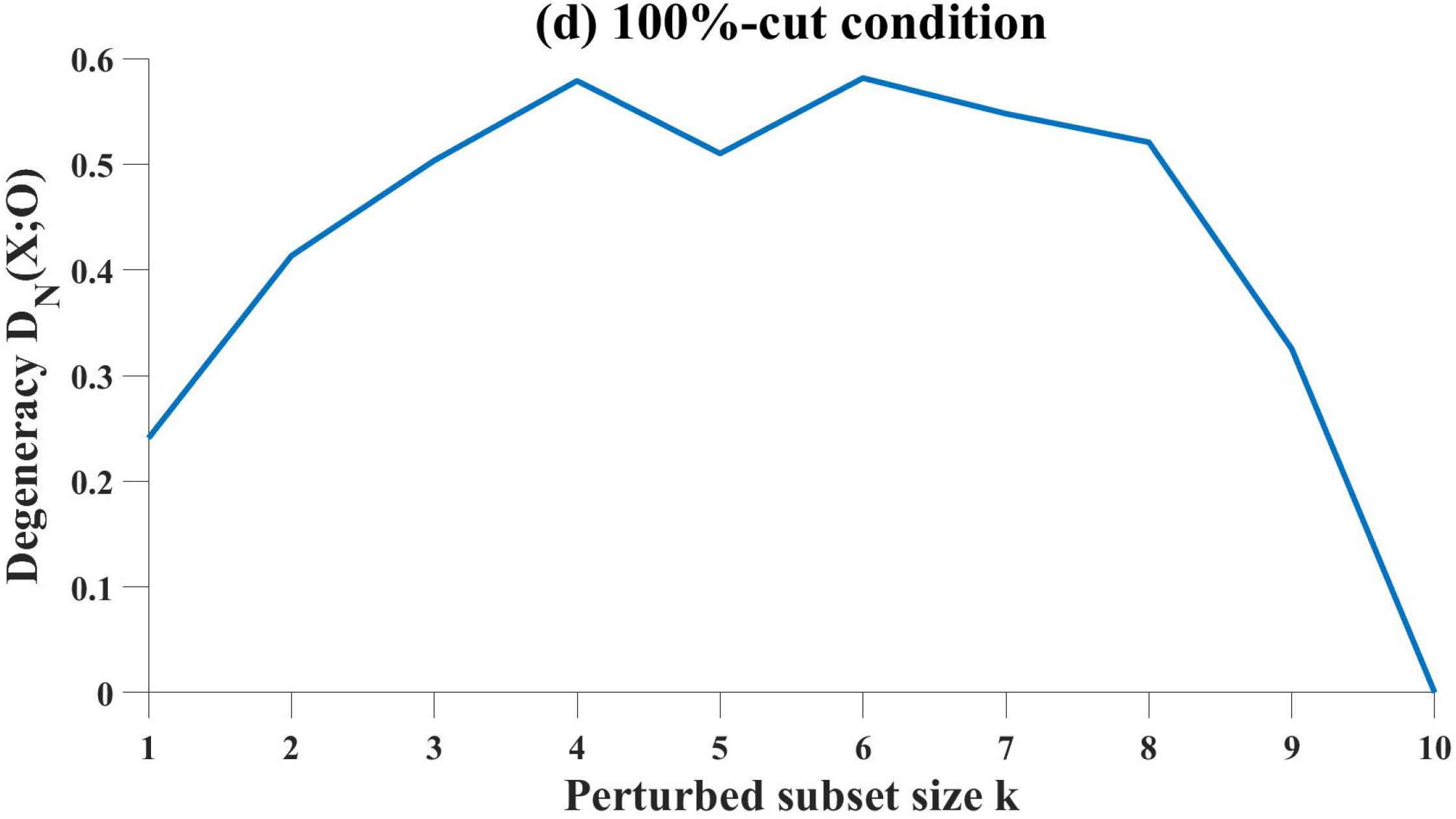
Degeneracy, (grey area) is computed (see Equation 3 in Methods) as the average MI between subsets of *X* and *O* under perturbation over increasing perturbed subset size *k*, in networks with (a, c) no lesions and with (b, d) 100%-cut condition.

In weighted networks with no connections, the average (overall-) MI shows a linear increase where degeneracy is zero [33]. Here, we have replicated this condition (i.e., 100%-of-edges-cut) in a similar way except, in our networks, nodes have preserved self-connections while all outgoing edges were lesioned. These networks also have initial variance due to the model (see details in Methods and Supporting Information) which is a system of stochastic differential equations (SDEs). Our results for no-edges-cut condition and the characteristic profile of degeneracy are comparable to corresponding findings in previous studies (Fig2 a, b).

Further inspection of partial degeneracy (see Methods for details) values from individual simulations showed that partial degeneracy can have a negative value for some conditions (only two of such conditions captured here for comparison, Figs 3a-b). Although we have observed that partial degeneracy is negative for different cut-conditions and different sizes of perturbed subset, when MI is averaged over the simulations for all the perturbed subset sizes k, <MI^P^(X^k^;O)>, degeneracy D_N_(X; O) was above zero in all conditions.

**Fig3.**
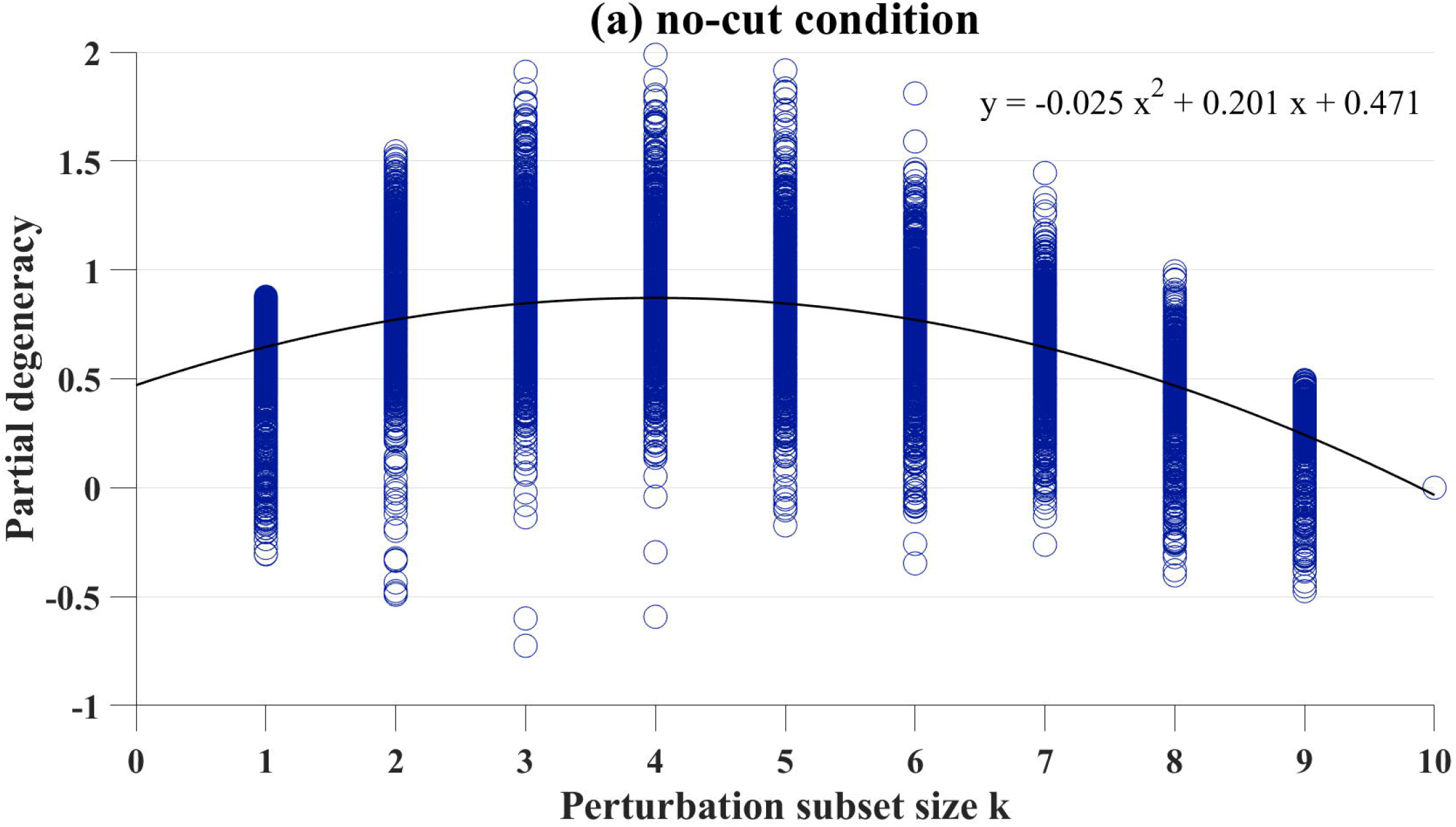
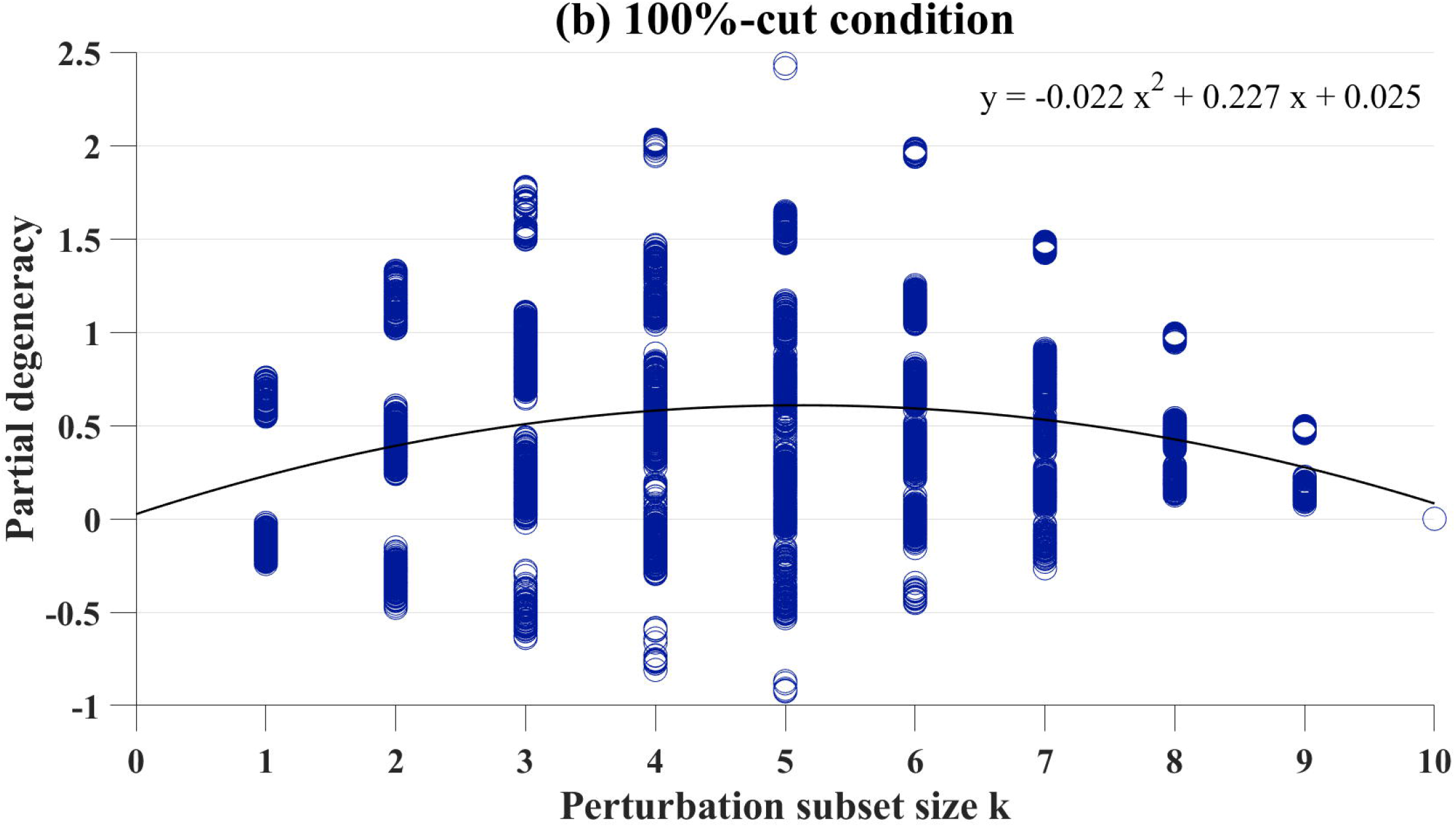
Partial degeneracy values from individual networks for each perturbed subset size k. Data from 10 (k) x 1000 simulations of networks with (a) no-cut condition and (b) 100%-cut condition. The distribution of partial degeneracy values for (b) shows clear modes in the data where there is an overlap of the perturbed subset and output sheet.

### Interactions of lesions and perturbations

Increases in both lesion extent and the number of nodes perturbed contribute to decreases in the degeneracy, raising the question of how these two factors may interact. We therefore conducted additional simulations in which each lesion condition (0%-100%, see Methods) was crossed with each perturbation condition (k = 1-10). Fig 4 (a-b) show how average partial degeneracy changes as a function of perturbation subset size k, given cut percentages. For both lesioning types, partial degeneracy peaks around when half of the nodes *(k ~ 5)* are perturbed in the system. The measure of degeneracy can detect existing isofunctionality between the different structures only when one of the structures is perturbed. When half of the nodes in the system are perturbed, we, thereby, maximize the probability of selecting/measuring the right structure for given degeneracy.

**Fig4.**
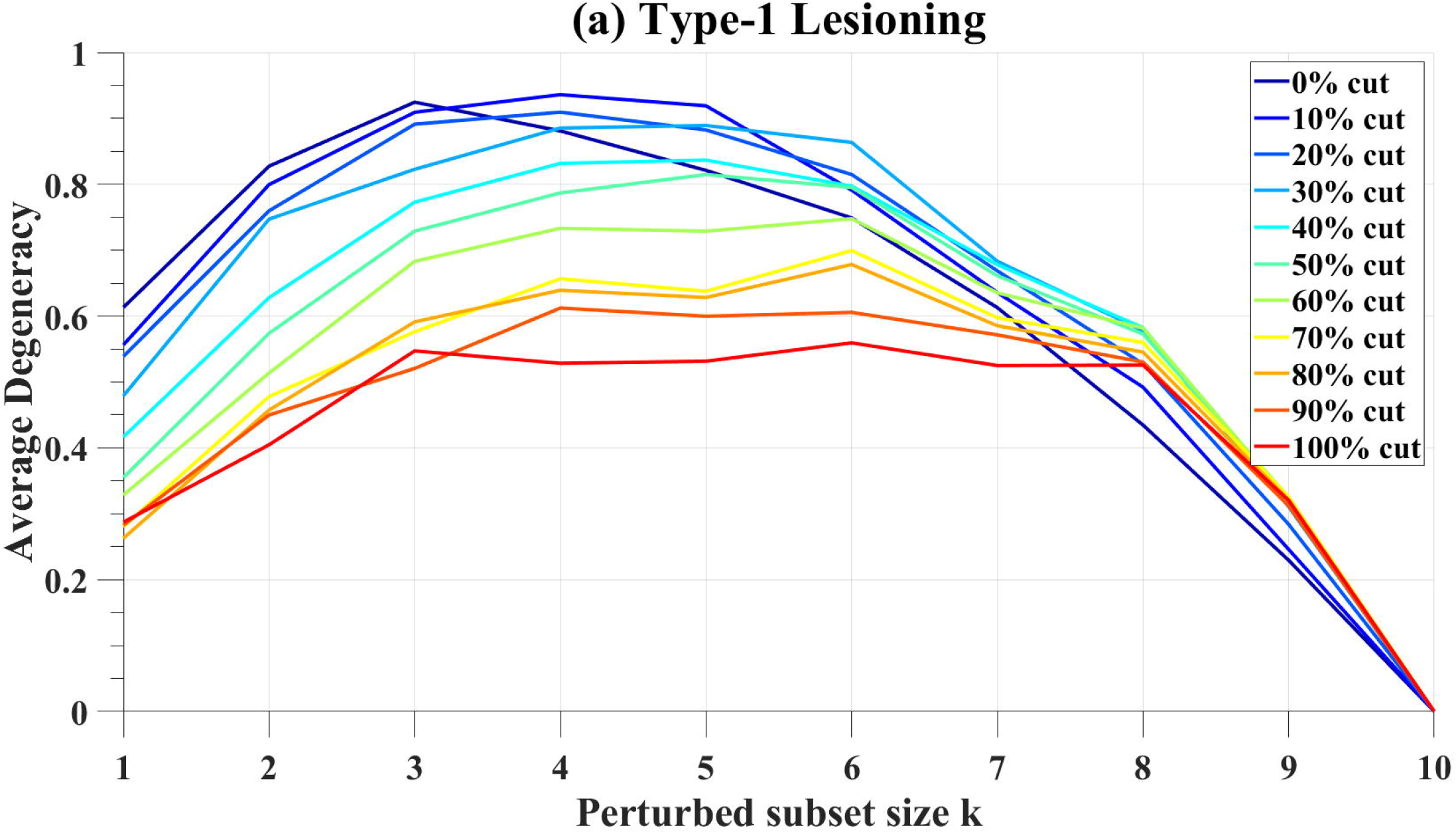
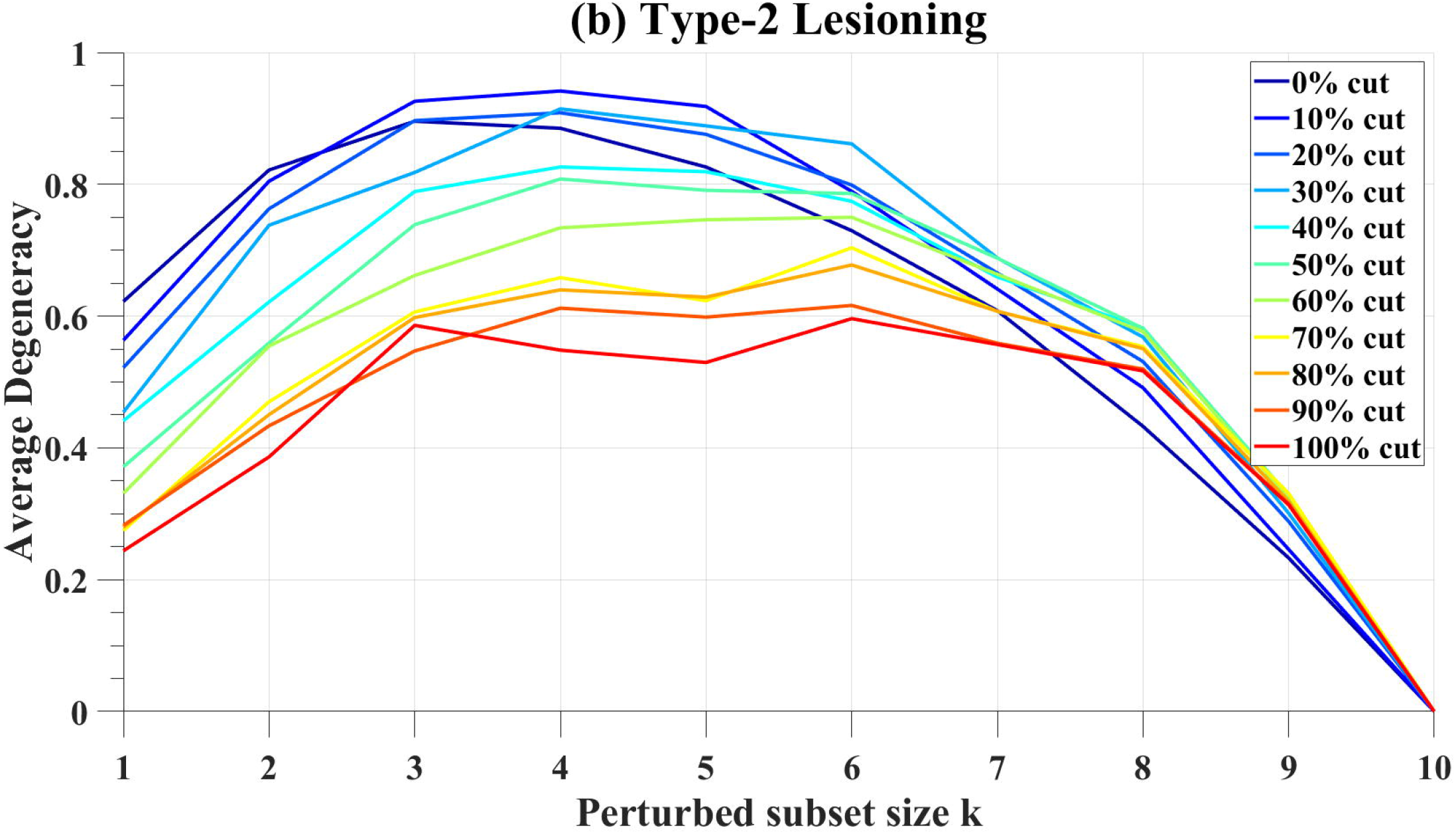
Average degeneracy computed as a function of perturbation subset size k in type-1 lesioning (a) and type-2 lesioning (b). Each line represents the cut condition for lesioned edges given the percentage of number of nodes in networks.

**Fig5.**
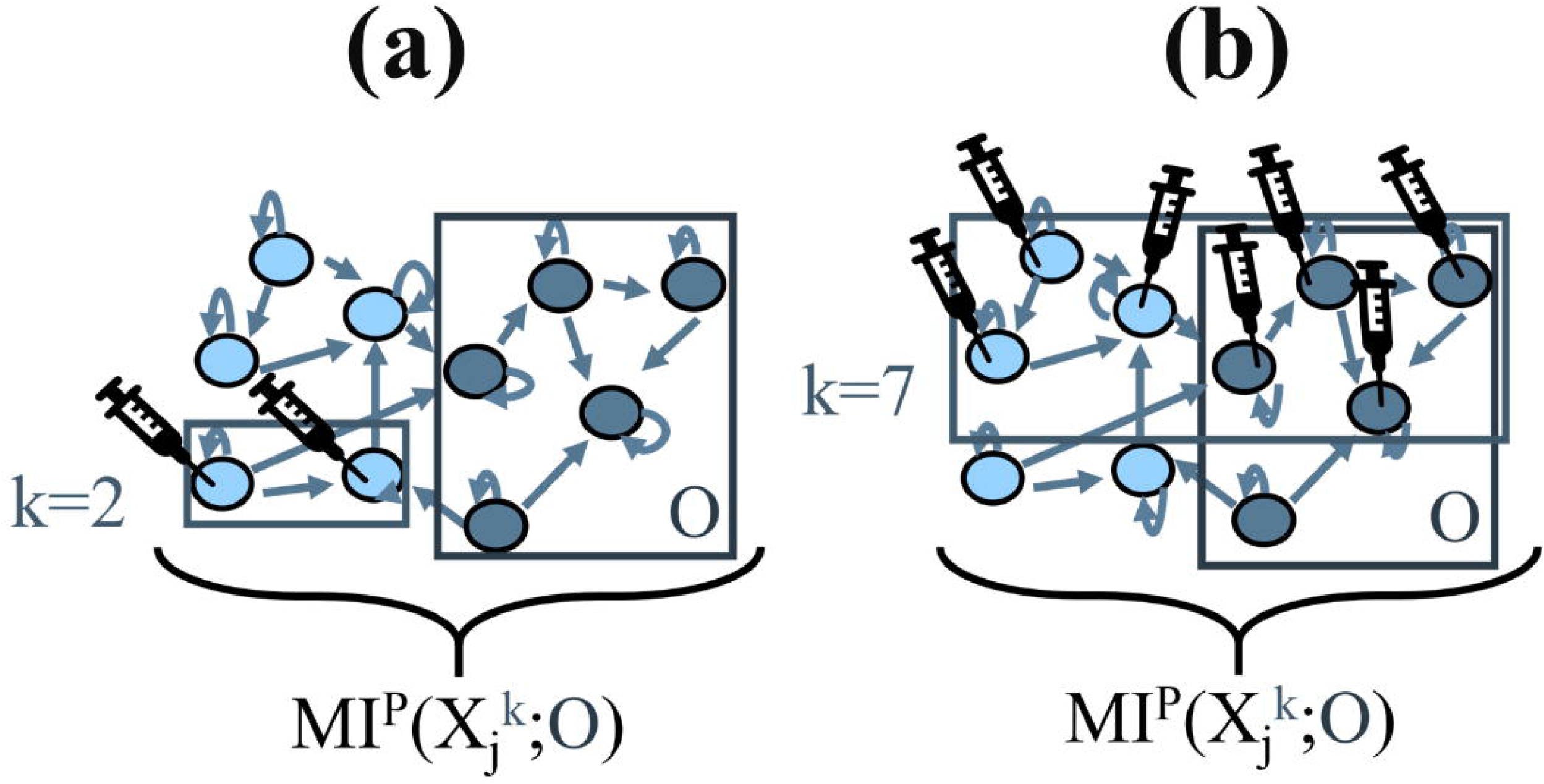
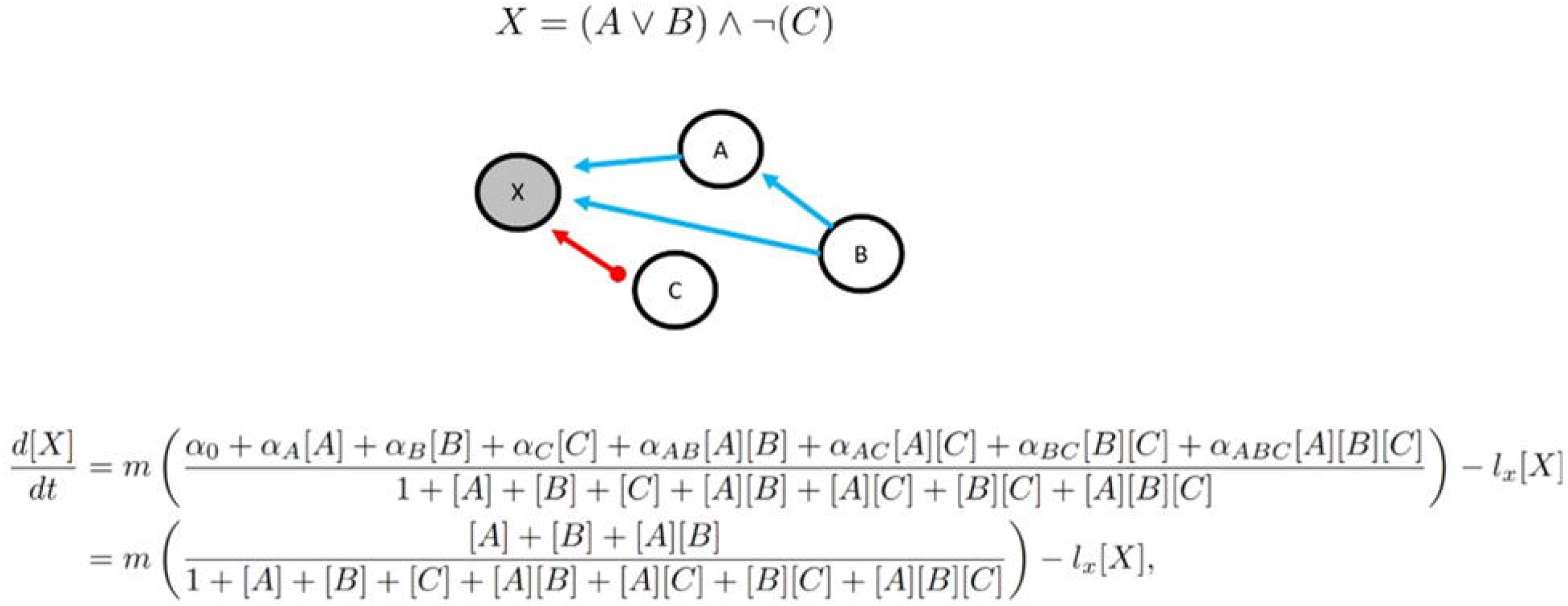
Illustration of RBNs under perturbation. A network of X, composed of nodes (light and dark blue circles, n = 10) that are interconnected. Arrows represent the edges for incoming, outgoing, and self-connections. Light blue circles represent randomly chosen perturbation subsets of nodes for k = 2 (a) and k = 7 (b). Perturbation (represented as syringes) of the nodes in boxes with k notation, is injected as a variance (uncorrelated noise). The box with O notation represents output sheet that is also consisted of randomly chosen set of (n/2 = 5) nodes (dark blue circles). For each network, MI is calculated for perturbed set size k and the output sheet O, for all subset sizes of perturbed set noted as j.

## Discussion

Although a variety of types of biological networks are thought to exhibit degeneracy, previous theoretical work has primarily focused on networks with weighted connections [33,40,59]. In this study, we demonstrate that degeneracy measures are also suitable for RBNs, frequently used to model gene regulation mechanisms. Although our simulations largely replicated previous studies investigating degeneracy in neurally-inspired networks, RBNs use Boolean logic operators rather than weighted connections to determine function. It therefore might have been the case that information-theoretic approaches developed for one class of networks might not have generalized correctly to a new class. By replicating previous findings using RBNs, we demonstrate that information-theoretic approaches are applicable to a broad range of network types.

Network and graph theoretic approaches, frequently formulated in terms of information theory, have been applied extensively to neuroscience [50–52,58,65–74] to predict individual differences, consequences of lesions, and ability to recover function following injury. Extending this approach to the study of GRNs opens the door to investigating the consequences of, and possible remedies for, genetic dysfunction. Because most genetic functions are performed by subsets of many components within functional modules [5,75], diseases may emerge due to disorganization of the components in these modules. Degeneracy measures can be recruited for predicting and inducing topological modifications (for example, ‘rewiring of diseased modules’ [75]) to achieve desired functional outcomes that have clinical significance, such as enhanced pharmaceuticals with better drug targets.

In addition to replicating previous results, we explored the impact of systematically manipulating network connectivity (lesioning) while decomposing degeneracy by size of the perturbed subset. In doing so, we identify a potential interaction between the number of perturbed nodes and the magnitude of the impact of lesions on network degeneracy. In networks in which ~20% of the connections are lesioned while 50% of the nodes are perturbed, it is observed that average partial degeneracy reaches its highest value among all other cut conditions and for all perturbed subset sizes. This can be interpreted as, for some conditions (here, k = 5 and ~20%-cut) might allow the expression of degenerate structures without compromising their function, whereas more lesioning would diminish both primary and degenerate structures, and more perturbation would confound the functions of the nodes. Likewise, if perturbation (of the number of nodes) is smaller, not all possible degenerate structures might have expressed in the network and, also, when the lesioning is less, degenerate structures might be unobservable since most of the primary structures are intact.

In the literature it has been shown that additional damage (gene/node deletions) can restore the function of previously compromised (metabolic) networks [76–78]. Here, we show that, up to a point, progressively lesioning a network results in increased degeneracy. This finding suggests that it might be possible to determine an optimum degree of lesioning and perturbation given a network to achieve higher degeneracy in the systems. Thus, partial degeneracy measures might be helpful to develop strategies to predict how to recover the function after damage.

As originally conceived, degeneracy was intended to capture the idea that identical functions could be carried out by distinct network structures. Intuitively, therefore, degeneracy would seem to have a lower bound at zero – in a network with no degeneracy, all structures would serve their own individual functions, and perturbation of those structures would disrupt network output related to the function served. Although on average degeneracy in our simulations tended to be above or equal to zero, we observed individual simulations in which partial degeneracy values were below zero. In previous studies ([40,59]), negative degeneracy has been observed especially for network models with increased biological fidelity.

One possible reason for the observation of negative degeneracy may be that the information-theoretic measure for degeneracy was originally developed for and tested on neural networks with no initial variance. As the biological fidelity of the models (thus, inherent variance in the systems) increases, for some conditions (lower coupling and lower connection probability [40] and networks with lower complexity [59]) negative degeneracy has been shown. However, we have not observed such effects on overall degeneracy measurements where 1000 MI values for each possible subset size k were averaged across random networks. Mathematically, degeneracy gets a negative value when the portion (*k/n*) of the MI between the whole system (*n*) and the output sheet (*n/2*) is higher than the average of the MI between the (perturbed) subset of the system (*k*) and the output sheet (*n/2*). However, the biological meaning/equivalence of negative degeneracy remains unclear. For studying more biologically realistic complex networks, adjustments in the tools for quantifying degeneracy may be needed.

## Methods

### Network Architecture

RBNs were initially proposed as simplified models for gene regulatory networks by Kauffmann [79] where network nodes represent the genes, and the edges represent the regulatory functions. A RBN constitutes a discrete dynamical system that has N nodes with K incoming edges (hence, also referred to as N-K models). Each node (gene) can be ON or OFF (1 or 0); a network of N binary nodes therefore has 2^N^ distinct states [79]. This system is state determined [79] according to Boolean functions that are assigned to each node randomly (from K^N^ possible states) where each node has a minimum of zero to a maximum of N inputs [80]. Such a state-space allows random network configurations which often leads to nonlinear dynamics. In this study, the nodes can have no inputs (but the self-connection) without an upper boundary, so that a gene can have a maximum of N inputs. The total number of nodes representing the genes, here, is *N = 10*, thus there are 2^10^ possible states. In RBNs, the state of the nodes in the network can be updated synchronously or asynchronously in discrete time steps. In this study, for simplicity purposes, a synchronous update rule is chosen.

RBNs have a well-defined function mapping scheme through logic (Boolean) operators which constitute the rules for connections that control the state of gene regulators. In our setting, operators *AND, NOT, OR* are randomly placed to generate rulesets (functions). If a node has only one input (that is one gene is connected to another gene) the probability of the function to have *NOT* operator is 0.5. If a node has two genes assigned to it (outgoing edges), the probability of these two genes to be connected to the node via *OR* operator or *AND NOT* operator is both 0.5. Outgoing edges are randomly distributed for each node with the condition that each node has at least one (thereby, connectivity is preserved) and at most two (more than one Boolean operator) edges mapped to the other nodes, while all the nodes have self-connections Thus, incoming edges are assigned to the nodes in a completely random fashion (allowing for the emergence of network hubs).

### Network Lesioning

Degeneracy, as a strategy (or design principle [10,21,81]) for networks to recover their function, refers to the rearrangement of (structurally different) components in a way that function/output remains the same even after a damage. In a network with high degeneracy, there are many possible network reconfigurations that can produce/recover the function. To test the potential factors that give rise to (higher/lower) degeneracy in networks, here, we induce interventions to the systems at the network-level by lesioning the edges.

Two different lesions were introduced to the synthetic networks. In type-1 lesioning, all outgoing edges from randomly chosen nodes were cut while the self-connections of the nodes and incoming edges were preserved. In the second type of lesioning, all incoming and outgoing edges were cut (except for self-connections) from randomly selected nodes given the percentage of total lesioned edges. In both types of lesioning, the edges are lesioned in increments of ten percent of the total number of the nodes given a network. For example, in type-1 30% cut condition, we have lesioned all the outgoing edges (except the self-connections) of the 3 randomly chosen nodes given a network of 10 nodes. Likewise, in type-2 30% cut condition, all the edges (incoming and outgoing, except the self-connections) of 3 randomly chosen nodes (out of 10 nodes total in a network) were lesioned. By 100%-cut condition (in both lesioning types), we refer to networks where no node is connected to the other, and the nodes have only self-connections.

For both lesioning types, 0%-cut condition refers to networks that are not lesioned, yet the edges are randomly disturbed (according to the method that is defined previously). This may lead some nodes to not have any edges (but the self-connection) due to random assignments of the rules, therefore, mimicking a (partially) lesioned condition.

### Biologically realistic random Boolean networks: Discrete to Continuous

Boolean functions describe how the states of the regulators control the state of the target genes [82]. In our study, Boolean functions are randomly generated for each simulation with incremental lesioning. To execute numerical simulations, we used the BoolODE pipeline by Pratapa et al. [82]. BoolODE systematically converts a random Boolean network into a system of SDEs that is a continuous model of gene regulation (for model specifications see SI Text 1 and SI Table). Time points in the numerical solution result in vectors of gene expression values that correspond to individual cells. That means for every analysis, each sampled time point is from a cell [82], and in this study, we sample from 990 time points (1000-10, first 10 timepoints treated as burn-in) for each gene in a single simulation and total of 1000 simulations are run for each lesioning percentage increment of 10s (from no cut condition to all 10 genes cut) which makes 10000 simulations for each type of lesioning and thus, 2 (lesioning type) x 10 (k subset of perturbed genes) x 11 (cut conditions, no-cut condition inclusive) x 10 synthetic cells with random gene regulatory mechanisms.

### Quantification of degeneracy in neural networks

To measure degeneracy, we used the mathematical framework described by Tononi et al. [33]. In this framework, degeneracy is characterized in terms of the average mutual information between subsets of elements within a system and an output sheet (which is also a subset of network *X*). The output sheet is a set of randomly chosen nodes in a network and its activity is a result of the interactions the other nodes in the system. Thus, activity in the output sheet represents the behavior or the response of the whole system.

From information theory, entropy (Shannon entropy with log base 2 for binary representation) is calculated from probability density functions for subsets of *X* (Equation 1). Then mutual information that measures the portion of entropy shared by the system subset 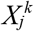 and the output *O*, is calculated as follows (*Equation 2*):

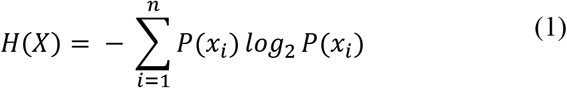

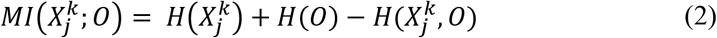

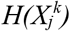 and *H(O)* are the entropies of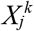 and *O* considered independently, whereas 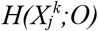 is the joint entropy of subset 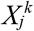 and output *O*. To measure degeneracy in the network, we need to determine the effects of the (subset of) element(s) on the entropy of the output - the behavior of the network. Since mutual information does not capture direction, however, mere calculation of mutual information is not enough to determine the contribution of the elements to the output of the system. To overcome this, perturbations (variance) are injected to the system. If no initial variance is assumed in the system, the value of mutual information between the network and the output is zero before any perturbation [33]. Variance (perturbation) is injected as uncorrelated random noise to each subset k.

Under such perturbations, mutual information of the system is computed as in *EQ3* and this procedure is repeated for all subsets of sizes *1 ≤ k ≤ n*. Then, degeneracy *D_N_ (X;O)* of *X* with respect to *O* can be calculated as:

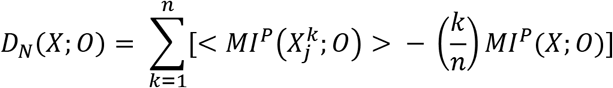

*MI^P^(X;O)* is MI for all elements to the output sheet, and <*MI^P^(X_j_^k^;O)*> is the average of the contribution of each perturbed subset size *k* to the output sheet.

### Quantification of degeneracy in random Boolean networks

Biological RBN simulations (via BoolODE) results in continuous unit activity in terms of gene expression vectors since the simulated networks are translated into nonlinear dynamical system (SI Text 1 and 2, SI Figure). To quantify degeneracy in RBNs we therefore discretize the gene expression vectors by taking the median activity for a unit. Activity that is above the median is set to 1, and activity below the median is set to 0.

We apply the degeneracy measures to the discretized gene expression vectors generated from the simulations. In our simulations, perturbations are systematically injected to the subset size k of genes as normally distributed (with *mean = 0*, and *standard deviation = 0.01*) random noise through the governing SDE. The number of elements (namely, the genes) is *n = 10* for all simulations with output sheet consisted of the activity of *O = n/2 = 5* elements, that is also randomized for each trial.

### Partial degeneracy in random networks

Degeneracy can be measured by alternate ways that are mathematically equivalent (see other definitions in [33]). The formal definition that we use in this study requires averaging over every MI measured between each node (unit, *j*) which are incrementally perturbed (from *k = 1* to *k = n*) and the output sheet for a given network structure (<*MI^P^(X_j_^k^;O)*>). However, in case where all the networks are randomly generated and the output sheet units are randomly chosen, an alternate way of computing <*MI^P^(X_j_^k^;O)*> is taking the average of MI measured for each random network that is perturbed once for a particular perturbed subset size k in range of *1 ≤ k ≤ n*. This way, degeneracy is measured for a specific subset given a network rather than for all possible subset sizes. Here, we call this measurement *partial degeneracy*.

## Supporting information

**S1 Text 1. Simulation platform.**

**S1 Text 2. Model specifications.**

**S1 Figure. A toy network with Boolean functions and translation into SDE.**

