## Supporting Information for "Degeneracy measures in biologically plausible random Boolean networks"

***Simulation Platform***

To generate synthetic gene expression data, we ran simulations on open source BoolODE pipeline (available from: <https://github.com/Murali-group/BoolODE>) V0.1 release. Following adjustments are made:

1) The BoolODE pipeline calls a text file that has a Boolean model as input, first, the model is translated into stochastic differential equations (SDEs) with added noise terms, then numerical simulations (that result in a stochastic time course) are run [82]. As input files, we have generated random Boolean models that are converted into text files and these models are called from BoolODE to generate datasets in the order described above.
2) Inherent noise parameter *c* is set to 0.3 (default value is 0) to introduce stochasticity.
3) In cases where simulations of network activity attenuates (goes to zero), after a trial with different initial conditions if the network still attenuates, these are assumed to be the result of that specific network configuration.

***Model Specifications***

Each random Boolean model is converted into an equivalent SDE that is a continuous model of gene regulation. The conversion steps are (using the same framework in GeneNetWeaver [82, 83]) as follows:

1) Each node in the network is assigned to a ‘gene’ variable representing the level of messenger RNA expression and a ‘protein’ variable representing the amount of transcription factors.
2) The amount of transcription factors is determined by a model that takes account of mRNA transcription and degradation rates, as well as protein translation and degradation rates. Transcription and translation are counterbalanced by the degradation of the mRNA and protein pools.
3) Transcription rate depends on the affinity of gene’s promoter to the transcription factor. The probability of each binding-configuration is computed. Then, the efficiency of transcription activation by a specific configuration of bound regulators is calculated (with added noise term to introduce stochasticity). The cooperative effects of regulator binding are set by the parameters of Hill threshold and Hill coefficient.

In this study, we have preserved default model specifications for gene regulation in V01 release of BoolODE pipeline. Perturbations are injected to genes (not to proteins) as Gaussian variance with *mean = 0* and *standard deviation = 0.01* at each time step of the simulations.

The gene regulatory model parameters are preserved as in the original mathematical model that is used in Pratapa et al. [82]. The parameters and their default values are as follows:

mRNA transcription rate (*m*) = 20

mRNA degradation rate (*l_x_*) = 10

Protein translation rate (*r*) = 10

Protein degradation rate (*l_p_*) = 1

Hill threshold (*k*) = 10

Hill coefficient (*n*) = 10

The states of each gene are determined by Boolean functions with input states representing the activity of the regulatory factors (incoming edges). All possible effects from incoming edges are converted into parameter α_p_ where “p” stands for the parent node (source of the edge). The value of the parameter α_p_ is the output of the Boolean function relating to the activity (/state, ON or OFF) of the target gene (child node) and the state of the regulatory factor (parent node).

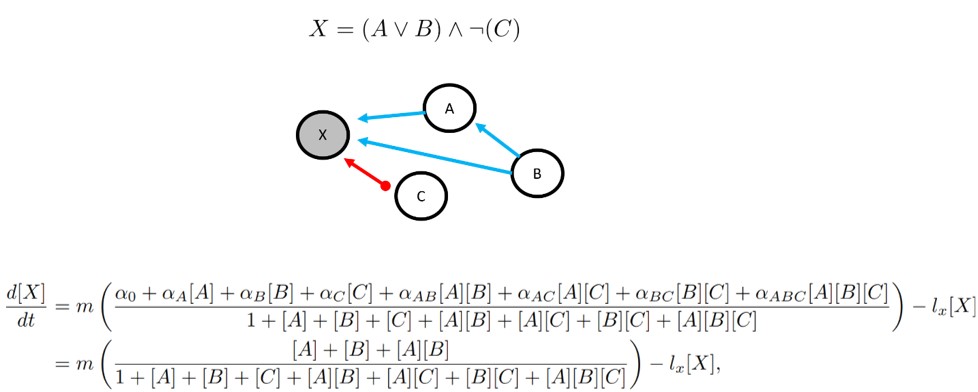

**Supporting Information Figure.** **A toy network with Boolean functions and translation into SDE.** The Boolean function (*top*) for the target gene X defined by the configuration of the incoming edges. Incoming edges show the regulatory control over the target gene, and are represented as arrows (*middle*), where upregulation correspond to blue arrows and red arrows represent inhibitory control. The equivalent SDE for gene X is retrieved from Pratapa et al. [82] (*bottom*).
